# Reducing the vicissitudes of heterologous prochiral substrate catalysis by alcohol dehydrogenases through machine learning algorithms

**DOI:** 10.1101/2023.09.26.559656

**Authors:** Arindam Ghatak, Anirudh P. Shanbhag, Santanu Datta

## Abstract

Alcohol dehydrogenases (ADHs) encompass three distinctive superfamilies: medium-chain dehydrogenase (MDRs), short-chain dehydrogenase (SDRs), and iron-containing alcohol dehydrogenases. MDRs and SDRs have pivotal roles in converting pro-pharmaceutical ketones into chiral alcohols, serving as active pharmaceutical intermediate catalysts in industries. Nevertheless, the pursuit of aligning enzymes to a one enzyme-one substrate paradigm is intricate and resource-intensive. This study undertook the catalysis of 284 antibacterial ketone intermediates, yielding distinct patterns for both MDRs and SDRs. We endeavored to leverage machine learning to discriminate between compounds subjected to the influence of these superfamilies. To this end, we curated a dataset comprising 33 features, encompassing 4 descriptors for each compound. Subsequently, an ensemble of machine learning techniques, including Partial Least Squares (PLS) regression, k-Nearest Neighbors (kNN) regression, and Support Vector Machine (SVM) regression, was harnessed. Furthermore, the assimilation of Principal Component Analysis (PCA) augmented precision and accuracy, refining compound classification and intensifying differentiation and demarcation within diverse compound classes. As such, this classification strategy emerged as a potent approach for discerning substrates amenable to diverse alcohol dehydrogenases, thereby mitigating the reliance on high-throughput screening in identifying the optimal enzyme for specific substrate preferences.

## Introduction

The world population is around 7.7 billion (and increasing), according to the 2019 revision of the United Nations population prospect report. In the next 30 years, it is predicted to grow by another two billion people, reaching over 11 billion by 2100 [1]. This has been fueled by an increase in the quality of living, change in fertility rate, migration and most importantly modern medicine. However, increased life expectancy does not imply that people are less unwell today than in the past, because good treatments are not available to everyone. As a result, discovering new medications and chemicals will continue to be critical for our survival. Additionally, chemicals for hygiene and cosmetics, food additives, fertilizers, insecticides, herbicides, plastics, nylon, rubber, paints, refrigerants, organometallics, polymers, liquid crystal displays in smartphones and televisions shall be required at an ever-increasing quantity.

Consequently, climate change is inevitable due to further industrialization. The UN intergovernmental panel predicts rise in global temperatures by 1.5°C in 2030. This level is deemed as dangerous, as it can destroy biodiversity and livelihood [2]. The current way of life comes at a significant cost in the form of toxic chemical waste due to reliance on petroleum-based goods for producing new chemicals. Many of them are manufactured under hard conditions like high temperatures and low pH [3]. Further, hazardous heavy metal reagents and toxic organic solvents are often used during the production. As a result, numerous manufactured chemicals and life-sustaining commodities products imperil the ecology and ecosystems of our world. Hence a sustainable method for producing active pharmaceutical ingredients (APIs) with less carbon footprint is essential. Consequently, enzymes are well explored sustainable catalysts for producing a APIs however, the path to obtain the perfect catalyst is often fraught with engineering obstacles as most of the APIs have non-natural origins.

In nature, metabolic processes take much longer to complete without enzymes, and this would consequently make living systems a distant reality. Enzymes help mitigate this difficulty and catalyze numerous chemical reactions in our bodies to support vital metabolic functions [4]. They interact with substrates through ‘active sites’ to stabilize the transition state and lower the activation energy required for synthesizing essential products. However, evolution doesn’t excel in making the best design rather, it only subsists to avoid selection pressure. With this thought, if an enzyme can fulfil conversion of a single substrate, then, it has certainly fulfilled its role if the reaction is deemed important for the survival of the organism. Hence, it is unsurprising that enzymes are specific and have a very narrow substrate range.

However, an industrial setup is a different ball game. Enzymes used in industrial processes, help in synthesis of pharmaceuticals, conversion of grain juices into lager and wine, the leavening of bread dough, manufacture of agrochemicals, synthetic flavors, biopolymers, waste remediation, and many others. Of these, bioremediation, and synthesis of active pharmaceutical ingredients (APIs) require enzymes to be versatile in converting multiple substrates [5]. The activity of APIs depend on the chirality of the molecules and the property of biospecificity is especially useful in synthesizing enantiopure chiral synthons. Therefore, the use of biocatalysts for manufacturing pharmaceuticals, has spurned interest for chiral intermediate production which help augment green synthetic methods of producing the final API [6]. This was made possible due to the advent of advancements in biocatalyst selection, screening, improvement, production, and supply technologies [7].

Chiral alcohols play a significant role in the activity of most APIs. They are well known for their stability and act as the building blocks for multiple drugs. In fact, more than 17% of the prescribed medicines are derived from these intermediates [8]. Some such examples of significant pharmaceutical intermediates include the production of Trusopt intermediate, which forms the Dorzolamide drug, which is useful in the reduction of ocular pressure among patients suffering from glaucoma and the Lipitor intermediate for Atorvastatin drug that prevents cardiovascular diseases as well is useful in the treatment of excess lipid in the body. Other examples of drugs include synthesis of optically active oxybutynin that is useful in the treatment of frequent urination, tomoxetine, denopamine, orphanedrine, tomoxetine, neobenodine and aprepitant, ezetimibe, floosool, ibrutinib, (R)-ladostigil, prosponine and sulopenem [9].

Similarly, drugs for the treatment of depression and Alzheimer’s disease such as fluoxetime and (R)-benzyl-4hyrdoxyl-2-pentynoate, respectively, are produced from chiral alcohol intermediates [10].

Haq *et al*. listed eight different APIs employed in pharmaceutical industry for production of varying chiral alcohols required for manufacture of drugs [11]. These include, 3,4- Methylenedioxyphenyl acetone for Talampanel, 1-N-carbobenzoxy-3-pyrrolidone for Barnidipine, 1-(3,5-Bis-trifluoromethyl-phenyl) ethanone for Aprepitant, 1-(2,6-Dichloro-3- fluro-phenyl)-ethanone for Crizotinib, 3-Trifluoromethyl acetophenone for MA-20565, 2-

Phenyl-1-thiazol-2-yl-ethanone 6 for Dolastatin, 1-(3-methoxyphenyl) ethenone for Rivastigmine and 3-Oxo-4- (2,4,5-trifluoro-phenyl) butyric acid methyl ester for Sitagliptin production. Apart from this, intermediates in the form of organophosphorus compounds has been useful in the synthesis of Tenofovir which is used for treating hepatitis-B and HIV, while Fosfomycin, Valinophos and Fosfazinomycin A, which are antibiotics and Alafosfalin, which is an antibacterial and antifungal drug [10].

Alcohol dehydrogenases (ADH) have attracted a lot of attention for their symmetry-breaking abilities in addition to their physiological responsibilities in metabolizing alcohols, aldehydes, and ketones [12]. As a result, they are prime candidates for producing chiral alcohols and hydroxyl compounds that are essential components in the synthesis of active pharmaceutical ingredients (API), which are used in the pharmaceutical business [6]. Recent advances in molecular biotechnology and structural bioinformatics have helped research and characterize enzymes at a faster pace. To this end, numerous alcohol dehydrogenases have been discovered in various organisms over the past few decades, beginning with *S. cerevisiae*, Horse liver, *Debaryomyces hansenii, Thermoanaerobium brockii, Rhodococcus erythropolis, Rhodococcus ruber, Pseudomonas* sp., *Lactobacillus brevis, Lactobacillus kefir* and *Candida parapsilosis* [13].

There are three types of alcohol dehydrogenases (ADH) viz. Type-I or Medium-chain dehydrogenase/reductase (MDRs) has 327–376 amino acid residues per chain. Many members use Zinc as cofactor [14]. Type-II dehydrogenases are short-chain dehydrogenase/reductase. There are two types viz. classical (250 residues long) and extended SDRs (∼350 residues long) [15]. Further, they don’t use any cofactor. Lastly, Type-III ADHs, comprise of iron-dependent alcohol dehydrogenases aka. FeADHs (385 residues long) [16]. Most characterized alcohol dehydrogenases are (S)-selective and adhere to the so-called Prelog rule. However, a structural and functional understanding of all three classes of enzymes is imperative for realizing the path forward with respect to green biochemistry of chiral alcohol synthesis. Upon comparison, Type I and Type III alcohol dehydrogenases have a narrow active site compared to the Type II enzymes or SDRs (Figure 1). Further, Type III enzymes are known to catalyze aliphatic primary and secondary alcohols. Therefore, for the production of chiral synthons production that predominantly consists aromatic rings it isn’t useful [17]. On the other hand many MDRs or Type I alcohol dehydrogenases are reported to produce chiral alcohols. Comparatively, SDRs have a larger substrate space and are quite diverse. They majorly encompass Classical and Extended SDRs where, the substrate space in the latter is larger and is known to catalyze multiple substrates [11][13].

**Figure 1.**
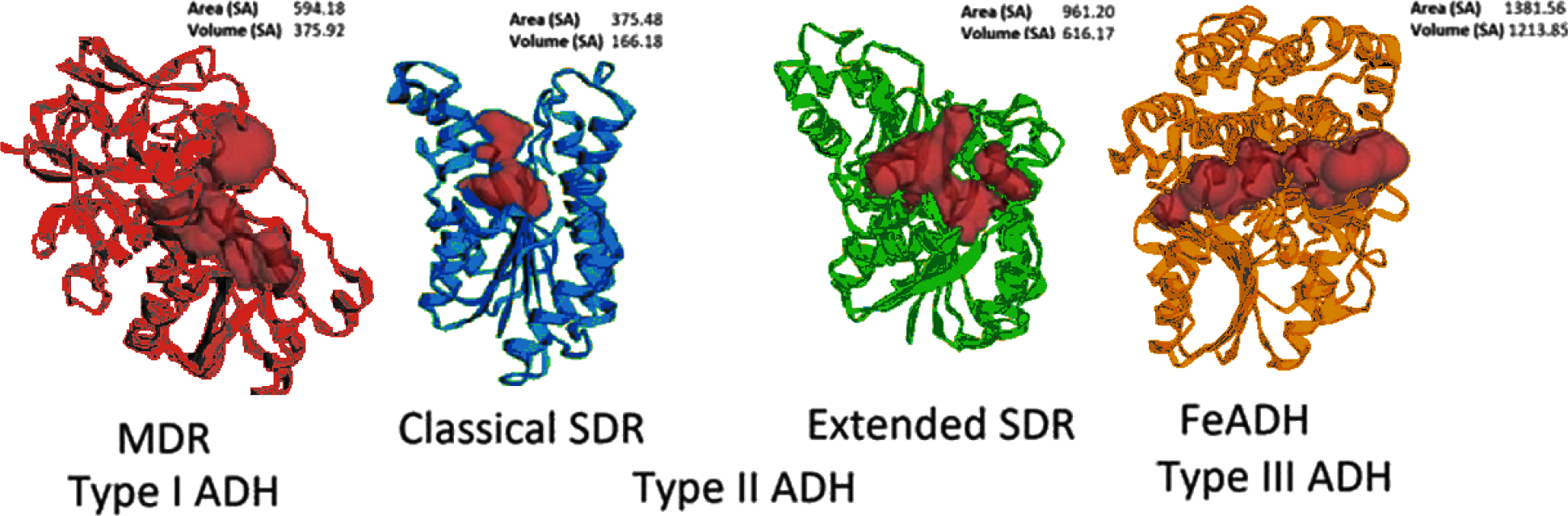
Substrate and cofactor binding spaces (maroon) of Type I (Medium-chain dehydrogenases/reductases or MDRs), Type II (Short-chain dehydrogenases/reductases or SDRs) and Type III (Iron alcohol dehydrogenases/reductases or FeADHs) with surface area and volume in Angstroms. The ratio between these characteristics denotes the ‘narrowness’ of Type III alcohol dehydrogenases (FeADHs) and ‘broadness’ of Extended SDRs.

## Results and Discussion

Before the advent of genome sequencing, ketoreduction was typically achieved by isolating organisms from rich physiological sources. Both Medium-Chain Dehydrogenases/Reductases (MDRs) and Short-Chain Dehydrogenases/Reductases (SDRs) play crucial roles in the detoxification of xenobiotics, dyes, and secondary metabolites in nature [18][19]. They have also been utilized in the degradation of dyes, pharmaceuticals, oil spills, and long-chain alkanes, among others [20]. Exploring these enzyme superfamilies has been facilitated by sourcing organisms from decaying wood, rotten fruit, wastewater, and hazardous landfills. However, the discovery and isolation of novel enzymes necessitate time-consuming and costly approaches such as genome and metagenome sequencing, high-throughput screening against expensive substrates, and more. Alternatively, enzyme engineering provides a viable solution for catalyzing specific substrates. Methods like directed evolution and site saturation mutagenesis involve substituting amino acids in the substrate binding site to identify the optimal combination for catalysis. Nevertheless, these techniques, including error-prone PCR and gene shuffling, require extensive screening processes, making them economically and temporally demanding [21].

Moreover, precise engineering is crucial, as even a minor change in atomic-scale structure can significantly impact catalytic performance. Although mesophilic enzymes are commonly employed, their stability and compatibility with other chemical steps are limited, necessitating the use of thermostable enzymes that can function at higher temperatures [22]. This is circumvented by using thermostable enzymes; however, they demonstrate flexibility at higher temperature and convert substrates at higher temperatures [23]. However, the energy and cost requirements for maintaining elevated temperatures during large-scale synthesis pose significant challenges. Consequently, identifying either promiscuous or specific enzymes for a given substrate using machine learning techniques presents an appealing prospect for industries.

The crux for identifying a desired substrate converting candidate lies in the turnover number of the enzyme. Hence, screening ADH types against a plethora of heterogeneous substrates and molding a paradigm to decipher catalytic pattern is thoroughly important. In the current study, we sought to find a pattern of catalysis among the ketoreductases belonging to different protein families. Therefore, we cloned and purified classical Short-chain dehydrogenases (SDRs) viz. FabG, ScoRed, ‘Extended SDRs’ DHK, ZRK and the MDR SSADH respectively (Figure 2). We chose 284 pro-chiral ketone having antibacterial property from eMolecules library to ascertain the substrate space of these enzymes. To this end, we determined the turnover number of the substrates by using them at saturating concentrations (1000 fold and 2000 fold) and subsequently comparing them using relative activity. Figure 3 shows the pattern of catalysis for the chosen enzymes.

**Figure 2.**
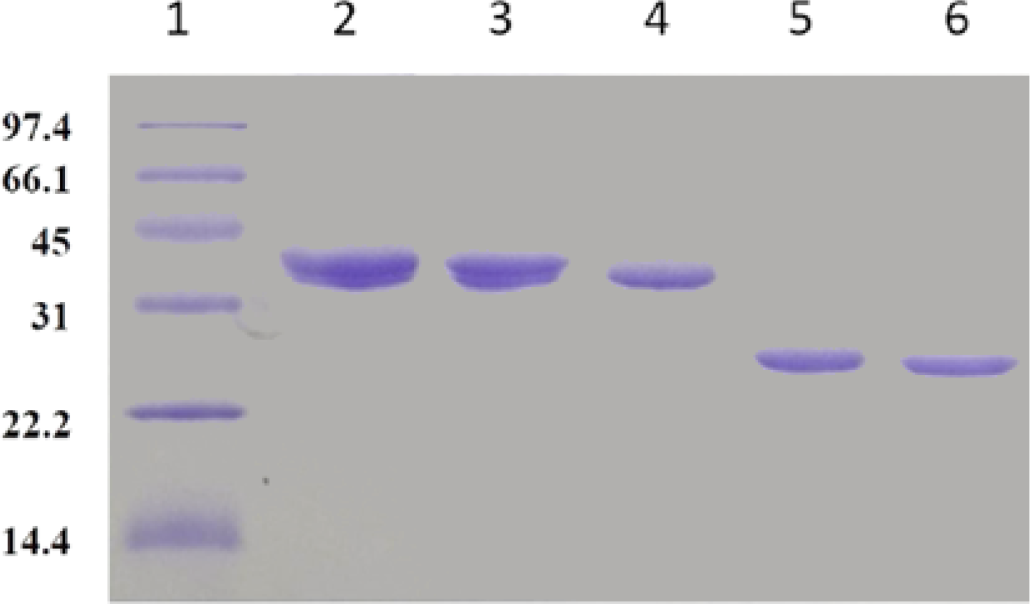
SDS-PAGE of reported SDRs and MDRs using Ni-NTA affinity chromatography. Lane 01: Low Range protein molecular weight marker, Lane 2: Lk-ADH from *Lactobacillus kefir* MDR Lane 3: Debaryomyces hansenii (DHK), Lane 4: SDR from Zygosaccharomyces rouxii (ZRK), Lane 5: *Streptomyces coelicolor* short-chain dehydrogenase reductase (ScoRed) and Lane 6: *Synechococcus* sp. PCC 4092 FabG.

**Figure 3.**
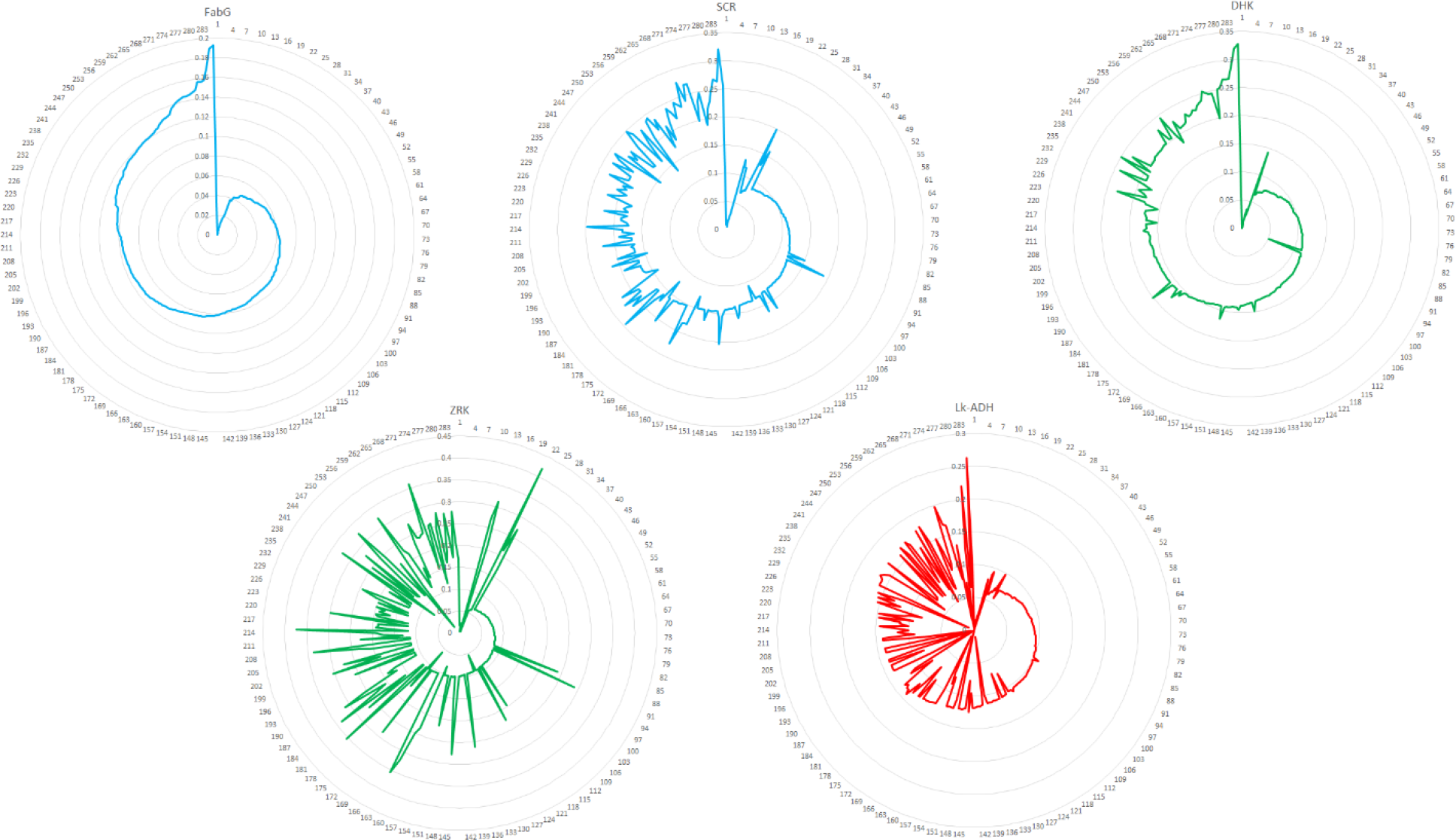
Web plot depicting the relative activity of the ketoreductases FabG, ScoRed, ZRK, DHK and LkADH against pro-chiral antibacterials. The classical and extended SDR activities are depicted in blue and green whereas, the MDR LkADH activity is depicted in red. A common catalytic pattern is observed between the SDR and MDR superfamily in the earlier ∼kp100 substrates. This clearly diverges for the later molecules thereby setting a pattern for exploration of substrates that can be catalyzed by both superfamilies.

Overall, the promiscuous ScoRed demonstrates spikes in relative activity compared to FabG thereby demonstrating the enzyme’s flexibility compared to FabG. The same is observed for DHK an Extended SDR. Ineterstingly, for ZRK (another reported promiscuous SDR) huge spikes are observed compared to other SDRs relative activity shows exvginations towards many substrates. Ineterstingly, for SSADH an MDR, the first 109 substrates have show similar catalytic activity as the SDR. However, there is a significant loss of activity (demonstrated as introversion in the plot) thereby, showing a clear preference of substrate in the superfamilies. In conclusion, MDRs and SDRs have common substrates for catalysis however, they diverge from each other where, Extended SDRs viz. DHK and ZRK show superior catalytic activity compared to their ‘classical’ counterparts in FabG and ScoRed.

Hence, a multidimensional data scaling approach was employed, considering only the compound structures, to create clusters. However, while Figure 4 demonstrates successful clustering for the initial 100-150 compounds, it fails to effectively cluster the remaining compounds. This indicates that relying solely on the chemical structure is insufficient for understanding enzyme activity. Further investigation is necessary to comprehend the intricacies of enzyme catalysis.

**Figure 4:**
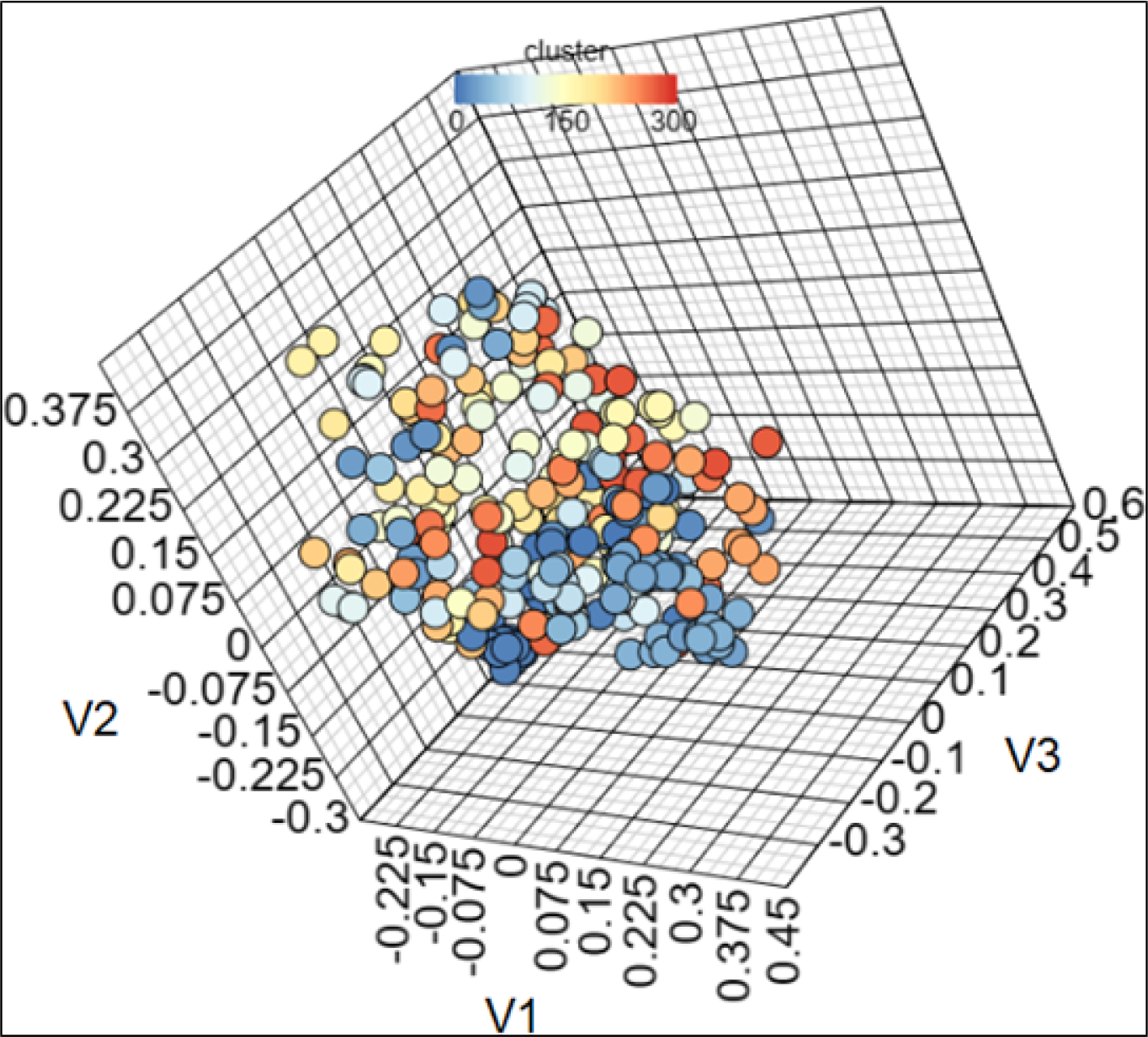
Multidimensional scaling clustering (MDS) of compounds at 0.7 cut-off showing pan-ADH catalyzed compounds (Blue) and compounds exclusively categorized by extended SDRs (White to orange). The clustering shows the former molecules grouped closer compared to the latter.

Unlike inhibitors, substrates have weak binding to enzymes, making it challenging to classify them compared to inhibitors or ligands for a specific protein. This limitation becomes apparent in a structure-activity landscape plot or SALI plot, where inhibitors of congeneric or non-congeneric series can be successfully clustered, but substrates with subtle associations with the protein of interest cannot. In fact, the analysis of nearest neighbors and the polar surface area of the compounds revealed a clear distinction between compounds that are reduced by SDRs (109-28) and MDRs (<109) (Figure 5A and 5B). However, this categorization is akin to placing substrates into two distinct groups without further insight. Therefore, additional algorithms incorporating various features and descriptors are needed to further classify the data and gain a more comprehensive understanding. Merely relying on compound structure is insufficient, and a more nuanced approach is required to accurately assess enzyme-substrate relationships.

**Figure 5:**
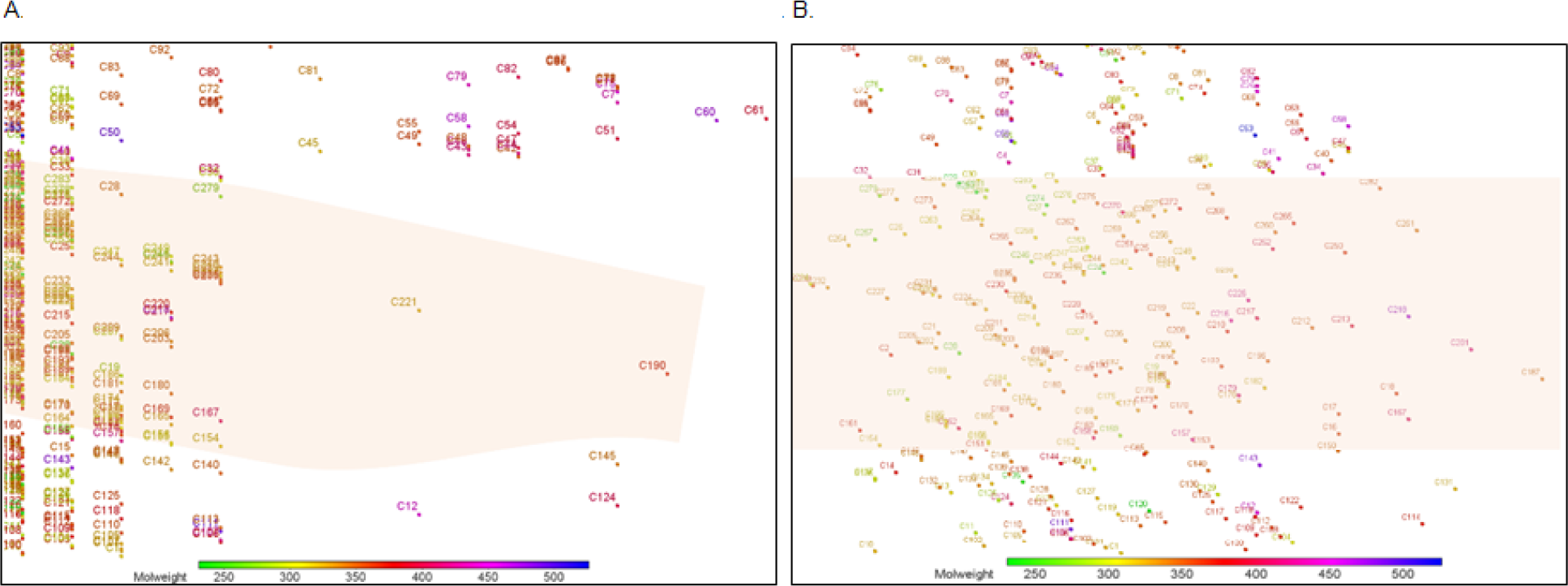
A Structure activity landscape (SALI) plot depicting the clustering of compounds using Nearest neighbor analysis (A) and Polar surface area (B) shows a clear demarcation between compounds which are reduced by SDRs and MDRs. The compounds that are catalyzed by Extended SDRs are highlighted in orange.

We utilized machine learning algorithms to effectively classify compounds based on their distinct features. The algorithms employed included Partial Least Squares (PLS) regression, k-Nearest Neighbors (kNN) regression, and Support Vector Machine (SVM) regression. Upon conducting regression analyses with the features against descriptors, we observed that the FragFP and SphereFP descriptors with PLS and kNN regression yielded the most favorable outcomes in terms of compound classification by using the R-square (∼0.8) value (Table 1). These regression models provided valuable insights into the relationships between the compound features and their respective classifications.

However, to further enhance the precision and accuracy of compound classification, we sought to employ Principal Component Analysis (PCA). It is a widely-used technique that facilitates dimensionality reduction while preserving the unique signatures and intrinsic characteristics of individual compounds. By employing PCA, we aimed to uncover the underlying patterns and variances within the compound dataset. By, integrating it into our classification pipeline, we reclassified the compounds using PLS regression, kNN regression, and SVM regression.

Intriguingly, this incorporation of PCA-driven classification led to notable improvements in the accuracy and reliability of compound categorization. It enhanced discrimination and separation between different compound classes, enabling more precise and successful classification outcomes (Figure 6). The compounds are divided into 9 baskets of which the first 4 are pan ADH catalyzed ketones whereas, the compounds in the other 5 are catalyzed by classical and extended SDRs. Overall, this inclusive approach involving machine learning algorithms, and PCA allowed us to achieve a more robust and refined classification of compounds.

**Figure 6:**
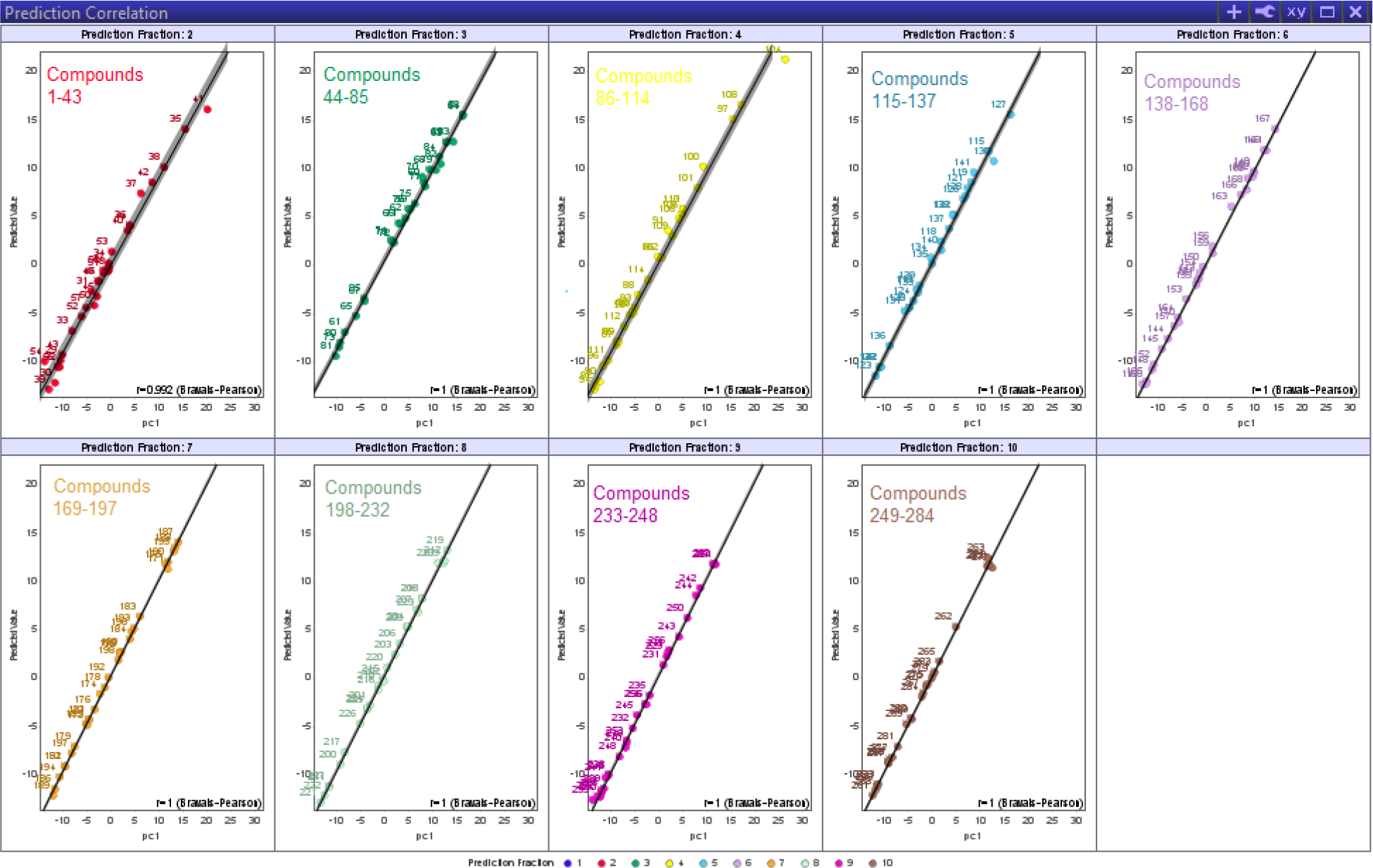
Figure depicting the Support Vector Machine (SVM) regression analysis of 284 antibacterials-ketones after using PCA analysis of 33 features. The r squared value is close to 1 thereby, demonstrating nearly seamless classification of substrates based on their ability to be catalyzed by different ADH types.

## Conclusions

In 2005, Swiss Industrial Biocatalysis Consortium (SIBC) analyzed the bio-catalytic needs of seven companies and indicated that the top need was for Ketoreductases that would reduce Prochiral ketones to chiral alcohols [24]. This activity is typically achieved by companies, such as Codexis Inc., Almac, Prozomix etc., by having a large library of enzymes from various sources [25][26][27]. The substrate is screened against the library to pick the best enzyme and if necessary, enzyme level engineering is conducted for improvement of catalytic efficiency. The strength of this strategy relies on availability of large, diverse enzyme libraries. However, with an ever-increasing number of synthetic substrates it would be useful to design an alternate strategy for finding the ideal catalyst. We propose the assessment of substrate range and its consistent pattern among various reported short and medium-chain dehydrogenases is a favorable gamble. To this end, a dataset of the chemical and structural features of the substrates is a useful asset. Although we have classified the compounds, the nuances related to extended SDRs such as ScoRed and ZRK where, some compounds are better catalyzed than others require more inspection. The current methods cannot help in this endeavor and require deep learning algorithms with more features to ascertain their ability to be catalyzed by ADHs. To this end, there are many chemical parameter calculators that can give one, two and three dimensional descriptors ranging >3000 [28]. Furthermore, using datasets for understanding substrate properties has been implemented in multiple studies [29][30][31]. Therefore, deducing this pattern using machine learning along with their corresponding turnover number can help predict novel substrate-enzyme pair for the designated API.

## Author contributions

**Santanu Datta:** Conceptualization, Investigation and supervision **Arindam Ghatak:** Writing, Methodology; formal analysis. **Anirudh P. Shanbhag:** Writing – review and editing; data curation.

## Acknowledgement

The assistance from Raksha Kini is gratefully acknowledged.

## Materials and methods

### I. Obtaining In vitro enzymatic data

#### (i) Purification of selected enzymes

Genes encoding the other heterogeneous enzymes namely, Sulfolobus solfataricus alcohol dehydrogenase (SSADH), Zygosaccharomyces rouxii SDR (ZRK), Hansenula polymorpha DL-1 peroxisomal 2,4-dienoyl-CoA reductase (Hketo), Synechococcus sp. PCC 7942 3-ketoacyl-[acyl-carrier-protein] reductase (FabG), Bacillus sp. ECU0013 ADH (ByueD) and Debaryomyces hansenii Ketoreductase (DHK) were codon optimized for E. coli and synthesized by Geneart (http://www.geneart.com).The genes were then sub-cloned into pET28a vector (Novagen) using the restriction sites NdeI and XhoI with an N-terminal His-tag in frame.

The recombinant proteins with an N-terminal hexa-histidine (His_6_) tag were purified by standard Ni-NTA affinity chromatographic method. Samples collected at every fraction were analyzed by SDS-PAGE (4-10%). A single, thickened band corresponding to a molecular weight of ∼39kDa was visualized in the final elution fraction for DHK and LkADH, ∼ 38kDa for *Zygosaccharomyces rouxii* SDR (ZRK), ∼26kDa for *Synechococcus sp*. PCC 4092 FabG, ∼27kDa for *Streptomyces coelicolor* short-chain dehydrogenase reductase (ScoRed) and ∼37kDa. Simultaneous amino acid sequence analysis was carried out to calculate the molecular weight of the protein. The proteins were desalted to remove the imidazole used during the purification process by a PD 10 column containing Sephadex G-25 Medium.

The buffers for protein storage and enzyme assays were as follows, 100mM sodium phosphate buffer, (pH 7.0) for FabG and ScoRed was 50mM Sodium phosphate buffer (pH 6.5) for ScoRed, 100mM Sodium Phosphate,100mM NaCl (pH7.4) for DHK and 50 mm MOPS buffer (pH 6.8) for ZRK and 10mM Sodium phosphate buffer with 300mM NaCl for LKADH.

#### (ii) Measurement of enzyme activity

In the enzyme assay, a concentration of approximately 1 μg/mL of the enzyme was employed for ketoreduction. The conversion of the substrate was determined using a spectrophotometer at a wavelength of 340 nm. To measure the kcat, saturating concentrations of substrates were utilized, which were approximately 500 times and 1000 times higher than the concentration of the enzyme. Consistent measurements of ketoreduction at both substrate concentrations indicated that the enzyme was operating at its maximum velocity. This information was utilized to calculate the kcat value, which is the ratio of the maximum velocity (Vmax) to the enzyme concentration (Et) in the reaction. A plot based on the kcat was then drawn using Microsoft excel for all the alcohol dehydrogenases.

### II. Initial clustering analysis

The initial analysis was performed using the ChemMine tool where, the clustering of compounds was performed using Muldimensional scaling (MDS) clustering. It entails using the smiles data from the chemicals and displaying a scatter plot based on the similarity of structures [28].

Further analysis was done using the SALI plot of the Data warrior software. Structure-Activity Landscape (SALI) Plot, also known as Structure-Activity Relationship (SAR) Landscape Plot, is a graphical representation used in medicinal chemistry and drug discovery to visualize the relationship between chemical structures and their corresponding biological activities. It provides a comprehensive view of the chemical space and activity space occupied by a set of compounds. In a SALI Plot, the chemical structures of the compounds are plotted on one axis, typically the x-axis, while the biological activities or properties of the compounds are plotted on the other axis, typically the y-axis. Each data point on the plot represents a specific compound and its corresponding activity value.

By examining the SALI Plot, patterns, trends, and clusters can be observed, indicating how structural modifications or variations influence the activity of the compounds. This helps in identifying regions of chemical space that are associated with desirable activities or properties. It also aids in the selection of compounds for further optimization or lead discovery in drug development.

### III. Datamining

The datamining of the chemicals was done using Datawarrior a popular opens source software which is used for clustering and classifying compounds based on the descriptors and features from chemical compounds [32].

#### (i) Descriptors used

The basis for understanding the nuances in the ketones catalyzed are ‘described’ in Data warrior tool by the aptly named ‘Descriptors’. The molecular structures of the molecules in a dataset are analyzed to extract specific molecular characteristics. These characteristics are combined to create a simplified description of the molecule, known as a descriptor. The simplest descriptor is a binary array where each bit represents the presence or absence of a particular characteristic in the molecule. These binary descriptors are also called fingerprints. More complex descriptors can take the form of a vector, a tree, or a simplified version of the original molecular structure.

If the goal is to filter a large collection of compounds based on their structural similarity, the default descriptor called FragFp is a suitable choice. It is readily available, requires minimal storage space, and similarity calculations are nearly instantaneous. For a more detailed assessment of similarity, such as distinguishing between stereo isomers or achieving optimal results in clustering or other similarity analyses, the SphereFP descriptor is recommended. Particularly, when creating an evolutionary library within an extensive virtual compound space, the SphereFP descriptor outperforms binary fingerprints in terms of quality. It takes into account the occurrence of multiple fragments and reduces the likelihood of hash collisions. The PathFp descriptor is a way to turn a small section of atoms in a molecule into a special code called a fingerprint. This fingerprint is made up of 512 numbers that are either 0 or 1. It helps us find and identify these small sections in the molecule. To make the fingerprint, we take each section and change it into a special text code that tells us what atoms are in the section and how they are connected. Then we change that code into a unique number. This number is used to set a specific position in the fingerprint to 1.

We used these descriptors as they suit our objective of finding out the structural differences between the compounds and also, the biochemical reasoning behind the catalysis of these compounds

#### (ii) Features selected for dataset

About 33 features were calculated for chemical compounds were calculated using the inbuilt Datawarrior tools for constructing the dataset:

Some well-known features such as molecular weight, molecular flexibility, molecular complexity, rotatable bonds, ring closures, aromatic atoms, stereocenters, sp3-atoms, aromatic rings and non aromatic rings, Hydrogen acceptors, Hydrogen donors, Monoisotopic mass, Electronegative atoms don’t need further description. However, some features that have been calculated in Datawarrior and play a major role in biocatalysis of compounds have been described below.

- Structure no.: The structures were assigned numbers corresponding to their susceptibility to reduction by the FabG enzyme. As all the enzymes follow a similar pattern, each compound was assigned a number, with 1 representing the least converted compound and 284 representing the most converted one.
- Monoisotopic mass: The theoretical monoisotopic mass of a molecule is calculated by adding together the precise masses of the most abundant naturally occurring stable isotope for each atom present in the molecule.
- cLogP: The clogP value provides information about the molecule’s ability to cross biological membranes and its potential for distribution within the body. As most of the molecules are antibacterial inhibitors from the eMoleules library we decided to use this descriptor.
- cLogS: It stands for the calculated logarithm of the aqueous solubility of a molecule. It is a predicted value that estimates the compound’s solubility in water. The cLogS value is derived using computational models or algorithms based on the molecule’s chemical structure and relevant molecular properties. It helps in understanding how soluble a compound is likely to be in aqueous solutions, which is extremely important for the determining the ability of enzyme to catalyse any given compound.
- Total surface area (Molecule TSA): The total surface area of a molecule refers to the sum of the areas of all the surfaces exposed by the molecule’s atoms. It represents the combined surface area that is available for interactions with other molecules or surfaces. The calculation of total surface area takes into account the shape and arrangement of the atoms in the molecule and provides insights into its physical properties and potential interactions with its environment.
- Relative PSA: It is a measure that quantifies the proportion of a molecule’s surface area that is polar in nature. It provides an estimation of the molecule’s polarity or hydrophilicity/hydrophobicity balance. RPSA is typically expressed as a percentage.
- Topological polar surface area: It is a measure that characterizes the polar nature of a molecule based on its topological features. It provides an estimation of the molecular surface area that is involved in polar interactions, such as hydrogen bonding. TPSA is calculated by summing the contributions of individual atoms or functional groups in the molecule, taking into account their connectivity and topological arrangement. The TPSA calculation considers specific atom types or functional groups that are known to contribute to the molecule’s polarity, such as oxygen or nitrogen atoms. TPSA is often used in drug discovery and medicinal chemistry to assess a molecule’s potential for interactions with polar regions of biological targets.
- Druglikeness: The druglikeness of a molecule refers to its potential to possess properties that make it suitable for further development as a drug candidate. It is a measure of how closely a molecule aligns with the desirable characteristics typically exhibited by known drugs
- Ligand efficiency: It is a measure used in drug discovery to assess the effectiveness of a ligand, typically a small molecule, in binding to its target protein or receptor. It evaluates the balance between the binding affinity of the ligand and its size or molecular weight.
- Lipophilic ligand efficiency (LLE) is a variant of ligand efficiency that specifically focuses on the balance between the binding affinity and the lipophilicity of a ligand. Lipophilicity refers to the molecule’s affinity for lipid or nonpolar environments.
- Ligand Efficiency lipophilic price (LELP) = logP / ligand efficiency as a useful function to depict the price of ligand efficiency paid in logP.
- Solvent excluded surface area (VDW surface): The solvent-excluded surface area (also known as the solvent-accessible surface area) is a related concept that represents the portion of the van der Waals surface that is inaccessible to a solvent molecule. It provides information about the exposed surface area of a molecule that can interact with other molecules or solvents.
- VDW volume: It refers to the volume occupied by a molecule based on its constituent atoms’ van der Waals radii. The van der Waals volume of an atom is an approximation of the space it occupies, considering the repulsive forces between atoms.
- Globularity SVD: Globularity, as determined through single value decomposition (SVD) of 3D atom coordinates, refers to a geometric property that quantifies the compactness or sphericity of a molecular structure. SVD is a mathematical technique used to analyze the properties and relationships of a matrix, in this case, the coordinates of the atoms in a molecule.
- Globularity volume (Globularity vol): In the context of molecular volume and surface analysis, it refers to a measure of how closely a molecule deviates from a perfect spherical shape. It is determined based on the comparison of the actual molecular volume and surface area to those of a sphere with an equivalent volume.

#### (iii) Machine learning

- kNN or k-Nearest neighbour regression: It is a supervised machine learning algorithm used for predicting continuous numeric values. It is a variant of the kNN algorithm, which is primarily used for classification tasks. In kNN regression, the algorithm determines the value of a new data point by considering its k nearest neighbors in the training dataset. The value of the new data point is predicted by calculating the average or weighted average of the target values of its k nearest neighbors.

To make a prediction using kNN regression, the algorithm calculates the distances between the new data point and all the training data points based on their feature values. The k nearest neighbors are selected based on the shortest distances. The target values of these nearest neighbors are then averaged to obtain the predicted value for the new data point. Here, the *k* value used was 3 and the distance used was Euclidean distance.

- SVM or Support Vector Machine regression: The algorithm aims to find a function that best fits the training data while minimizing the prediction errors. It works by constructing a hyperplane that maximizes the margin between the predicted values and the actual data points. The goal is to find the hyperplane that provides the best balance between fitting the training data and generalizing to new, unseen data. In SVM regression, data points that fall within a certain margin around the hyperplane are considered support vectors and play a crucial role in determining the regression function. The algorithm attempts to find the optimal regression function by identifying the support vectors that contribute the most to the regression performance.

One key parameter in SVM regression is the regularization parameter, often denoted as C. It controls the trade-off between achieving a small training error and minimizing the complexity of the regression function. A higher value of C allows for a smaller margin but potentially better fitting of the training data, while a lower value of C encourages a larger margin and better generalization to new data. The following parameters were used for classification viz. SVM Regression Type=3 Gamma=0.0 Epsilon=5.0 Kernel=2 C=130.0

- PCA, or Principal Component Analysis: We used Datawarrior to conduct principle component analysis after selection of features. It is a technique used to simplify and understand complex data. It helps us find the most important patterns and relationships in the data. These principal components are like new ways of looking at the data that capture the most important differences between the individuals. By using PCA, we can reduce the complexity of the data, visualize it in a simpler way, and uncover the main patterns and trends that might be hidden in the original measurements. We performed PCA for all the aforementioned descriptors and used kNN, PLS and SVM algorithm to classify the data.

Our objective was to evaluate multiple algorithms using different features derived from descriptors and assess their R squared values. The R squared value ranges between 0 and 1, where 0 indicates no correlation and 1 signifies a strong association or classification of the data. By employing various algorithms on the calculated features, we aimed to determine the level of correlation and classification achieved.

