## Supplementary Table 1 for "Reducing the vicissitudes of heterologous prochiral substrate catalysis by alcohol dehydrogenases through machine learning algorithms"

| Descriptor | PathFP | | | SphereFP | | | FragFP | | | SkelSpheres | | |
|---|---|---|---|---|---|---|---|---|---|---|---|---|
| **Algorithm** | **kNN** | **PLS** | **SVM** | **kNN** | **PLS** | **SVM** | **kNN** | **PLS** | **SVM** | **kNN** | **PLS** | **SVM** |
| Aromatic rings | 0.013 | 0.65 | 0.64 | 0.48 | 0.78 | 0.77 | 0.83 | 0.8 | 0.79 | 0.26 | 0.23 | 0.26 |
| Aromatic atoms | 0.014 | 0.54 | 0.77 | 0.46 | 0.8 | 0.68 | 0.79 | 0.85 | 0.81 | 0.3 | 0.27 | 0.3 |
| Non aromatic rings | 0.023 | 0.73 | 0.62 | 0.53 | 0.79 | 0.8 | 0.83 | 0.86 | 0.87 | 0.31 | 0.29 | 0.31 |
| cLogP | 0.03 | 0.43 | 0.78 | 0.57 | 0.79 | 0.77 | 0.82 | 0.86 | 0.85 | 0.29 | 0.3 | 0.29 |
| cLogS | 0.16 | 0.8 | 0.81 | 0.35 | 0.8 | 0.81 | 0.8 | 0.83 | 0.83 | 0.28 | 0.31 | 0.28 |
| Monoisotopic mass | 0.013 | 0.56 | 0.71 | 0.43 | 0.79 | 0.81 | 0.84 | 0.88 | 0.85 | 0.29 | 0.37 | 0.29 |
| Molecular flexibility | 0.02 | 0.8 | 0.77 | 0.44 | 0.8 | 0.7 | 0.83 | 0.89 | 0.78 | 0.32 | 0.42 | 0.32 |
| Stereocenters | 0.013 | 0.82 | 0.68 | 0.46 | 0.77 | 0.81 | 0.79 | 0.81 | 0.83 | 0.31 | 0.41 | 0.31 |
| Symmetric atoms | 0.01 | 0.79 | 0.76 | 0.55 | 0.77 | 0.73 | 0.81 | 0.79 | 0.78 | 0.31 | 0.32 | 0.31 |
| Total molecular weight | 0.05 | 0.81 | 0.79 | 0.45 | 0.77 | 0.73 | 0.82 | 0.78 | 0.79 | 0.31 | 0.35 | 0.31 |
| Globularity volume | 0.06 | 0.82 | 0.63 | 0.34 | 0.8 | 0.69 | 0.76 | 0.89 | 0.82 | 0.29 | 0.41 | 0.29 |
| Molecular complexity | 0.13 | 0.79 | 0.82 | 0.52 | 0.8 | 0.68 | 0.78 | 0.81 | 0.87 | 0.29 | 0.3 | 0.29 |
| Globularity VWD | 0.03 | 0.82 | 0.82 | 0.56 | 0.8 | 0.85 | 0.82 | 0.79 | 0.89 | 0.24 | 0.31 | 0.24 |
| Polar Surace area (PSA) | 0.02 | 0.84 | 0.82 | 0.37 | 0.78 | 0.78 | 0.77 | 0.84 | 0.83 | 0.26 | 0.34 | 0.26 |

| | | | | | | | | | | | | |
|---|---|---|---|---|---|---|---|---|---|---|---|---|
| Relative PSA | 0.04 | 0.65 | 0.8 | 0.4 | 0.77 | 0.78 | 0.83 | 0.78 | 0.78 | 0.25 | 0.4 | 0.25 |
| Rotatable bonds | 0.14 | 0.67 | 0.76 | 0.48 | 0.78 | 0.86 | 0.83 | 0.84 | 0.83 | 0.28 | 0.41 | 0.28 |
| Shape Index | 0.01 | 0.72 | 0.65 | 0.34 | 0.78 | 0.7 | 0.77 | 0.85 | 0.82 | 0.32 | 0.24 | 0.32 |
| Symmetric atoms | 0.04 | 0.45 | 0.72 | 0.49 | 0.79 | 0.72 | 0.76 | 0.84 | 0.81 | 0.32 | 0.27 | 0.32 |
| VDW volume | 0.02 | 0.43 | 0.75 | 0.44 | 0.78 | 0.9 | 0.84 | 0.87 | 0.89 | 0.32 | 0.37 | 0.32 |
| VDW surface | 0.02 | 0.42 | 0.79 | 0.48 | 0.78 | 0.89 | 0.75 | 0.79 | 0.79 | 0.29 | 0.39 | 0.29 |
| Principle cluster of all features | 0.92 | 0.89 | 0.96 | 0.99 | 0.93 | 0.94 | 0.94 | 0.95 | 0.95 | 0.49 | 0.42 | 0.36 |
